# Failure to resolve inflammation contributes to juvenile onset cardiac damage in a mouse model of Duchenne Muscular Dystrophy

**DOI:** 10.1101/2024.08.15.607998

**Authors:** James S. Novak, Amy Lischin, Prech Uapinyoying, Ravi Hindupur, Young Jae Moon, Surajit Bhattacharya, Sarah Tiufekchiev, Victoria Barone, Davi A. G. Mázala, Iteoluwakishi H. Gamu, Gabriela Walters, Jyoti K. Jaiswal

**Affiliations:** Center for Genetic Medicine Research, Children’s National Research Institute, Children’s National Research and Innovation Campus, Children’s National Hospital, Washington, D.C., 20012, USA; Departments of Pediatrics and Genomics and Precision Medicine, The George Washington University School of Medicine and Health Sciences, Washington, D.C., 20037, USA; Columbian College of Arts and Sciences, The George Washington University, Washington, D.C. 20052, USA; Neuromuscular and Neurogenetic Disorders of Childhood Section, National Institute of Neurological Disorders and Stroke, National Institutes of Health, Bethesda, MD, USA; Department of Biochemistry and Orthopaedic Surgery, Jeonbuk National University Medical School and Hospital, Jeonju, 54907, Republic of Korea; Integrated Biomedical Sciences, The George Washington University School of Medicine and Health Sciences, Washington, D.C., 20037, USA; Department of Kinesiology, College of Health Professions, Towson University, Towson, MD, 21252, USA

**Keywords:** Duchenne muscular dystrophy, cardiomyopathy, pathology, D2-mdx, B10-mdx, inflammation, inflammatory response, pro-resolution pharmacology, macrophage, macrophage migration, neutrophil, cytokine signaling, extracellular matrix remodeling, fibrosis, calcification

## Abstract

Absence of dystrophin protein causes cardiac dysfunction in patients with Duchenne muscular dystrophy (DMD). Unlike boys with DMD, the common mouse model of DMD (B10-*mdx*) does not manifest cardiac deficits until late adulthood. This has limited our understanding of the mechanism and therapeutic approaches to target the pediatric onset of cardiac pathology in DMD. Here we show that the *mdx* mouse model on the DBA/2J genetic background (D2-*mdx*) displays juvenile-onset cardiac degeneration. Molecular and histological analysis revealed that cardiac damage in this model is linked to increased leukocyte chemotactic signaling and an inability to resolve inflammation. These deficiencies result in chronic inflammation and fibrotic conversion of the extracellular matrix (ECM) in the juvenile D2-mdx heart. To address these pathologies, we tested the utility of pro-resolution therapy to clear chronic cardiac inflammation. Use of an N-formyl peptide receptor (FPR) agonist helped physiologically resolve inflammation and mitigate the downstream events that lead to fibrotic degeneration of cardiomyocytes, preventing juvenile onset cardiac muscle loss. These results establish the utility of D2-*mdx* model to study events associated with pediatric-onset cardiac damage and demonstrates pro-resolution therapy as an alternate to anti-inflammatory therapy for treating cardiac degenerative pathology responsible for cardiomyopathy in DMD patients.

## Introduction

Duchenne Muscular Dystrophy (DMD) is a severe and progressive muscle disease caused by the absence of dystrophin protein^1, 2, 3, 4^. Dystrophin plays a crucial role in maintaining the integrity of the sarcolemmal membrane by facilitating the assembly and function of the dystrophin-associated protein complex, thus its absence renders skeletal muscle cells more susceptible to mechanical damage, and increased muscle degeneration^5, 6, 7^. Dystrophin deficiency in cardiomyocytes also increases their vulnerability to sarcolemmal damage, cell death, chronic inflammation and cardiac fibrosis. These pathologies also manifest in patients and lead to the thinning of the left ventricle (LV) wall, causing their progressive dilation and dilated cardiomyopathy that results in heart failure^4, 8^.

Patients with DMD experience symptoms early in life, with cardiac deficits being a major contributor to premature mortality not only in DMD, but also in DMD carriers and in Becker Muscular Dystrophy (BMD)^9, 10, 11^. While the common B10-*mdx* model exhibits skeletal myopathy at an early age, the cardiac deficit is not evident until late adulthood^12, 13^. The advent of the D2-*mdx* model identified greater disease severity and fibrosis as compared to B10-*mdx* even in younger mice^14, 15, 16, 17^. Excess skeletal muscle fibrosis in the D2-*mdx* model, results from an increase in transforming growth factor beta (TGFβ) signaling, and cardiac deficit is reported as early as adulthood (16-weeks of age)^17, 18, 19^.

While therapeutic approaches to address cardiac deficit in patients with dystrophin deficiency are a topic of active investigation, anti-inflammatory glucocorticoids (GCs), angiotensin-converting enzyme inhibitors (ACEi), and angiotensin receptor blockers (ARBs), are commonly used for these patients^11, 20, 21, 22^. Anti-inflammatory therapy by GC has the longest history of use in DMD patients and has mixed reports of cardiac benefit and side effect in both DMD patients and mouse models^23, 24, 25, 26, 27^. An anti-inflammatory protein activated by GCs that mediates the GC efficacy is Annexin A1 (AnxA1)^28, 29^. Independent of its widely known role in membrane fusion, AnxA1 is required for muscle repair by regulating inflammation by way of Formyl Peptide Receptor (FPR)-mediated signaling^30, 31^. In this regard, AnxA1 works similarly to the endogenous pro-resolving lipid mediators Lipoxin A4, and Resolvin D1 by helping resolve acute inflammation by binding the FPRs^29, 32^. This feature of AnxA1 has led to the advent of natural and synthetic agonists of FPR2, that unlike GCs, reduce inflammation by promoting resolution of inflammation instead of suppressing the tissue’s inflammatory response^33^.

Use of FPR2-agonists reduces acute cardiac damage in various tissue injury models including myocardial infarction, where it restricts premature heart failure and restores tissue function^34, 35, 36, 37, 38^. Just as endogenous FPR2 agonists, a nanomolar dose of a synthetic agonist BMS-986235 also resolves chronic inflammation in preclinical models, which has led to its progress to clinical studies^38^ (Clinical Trial NCT03335553). This drug activates macrophage transition to a pro-resolving (M2-like) state by enhancing phagocytosis and neutrophil apoptosis, that regulates their chemotaxis, all of which help improve mouse survival, reduce scarring, and preserve tissue^38, 39^. These are desirable features of therapies to target cardiac inflammation and fibrosis associated with the cardiac pathology observed in DMD patients.

To understand the early onset of cardiac dysfunction in the D2-*mdx* model, we investigated the factors that distinguish the pediatric initiation of cardiac dysfunction as compared with the late adult onset in B10-*mdx* model. This revealed onset of cardiac pathology in juvenile (< 6-weeks old) D2-*mdx* mice, and showed this is associated with excessive immune infiltration, fibrotic ECM replacement, and degenerative remodeling of cardiac ventricular walls. It identified increased leukocyte chemotactic signaling and failure to resolve inflammation as major contributors to the initiation of cardiac pathology. To address this deficit, we test a drug-based pro-resolution therapy for a preclinical evaluation as a therapeutic approach to mitigate early-onset cardiac damage in DMD mouse models and patients.

## Methods

### Animals

All animal procedures were conducted following guidelines for the care and use of laboratory animals as approved by the Institutional Animal Care and Use Committee (IACUC) of Children’s National Research Institute (CNRI). The C57BL/10ScSn-DMD^mdx^/J (B10-*mdx*) and D2.B10-DMD^mdx^/J mouse models of DMD were utilized for all experiments and both harbor the same nonsense point mutation in exon 23 of the dystrophin (*Dmd*) gene thereby abolishing dystrophin protein expression ^40, 41^. The C57BL/10ScSn/J (B10-WT) and DBA2/J (D2-WT) mouse models were used as age-matched, model-specific controls. Mice were obtained from the Jackson Laboratory and were housed at the CNRI Comparative Medicine Unit where they were provided daily monitoring, food, water and enrichment ad libitum, while being maintained under 12 h light/dark cycles.

### Tissue harvesting and sample collection

Mice were euthanized via CO_2_ inhalation and cervical dislocation at designated ages corresponding to specific stages of disease progression. Muscles were surgically removed, mounted on cork with tragacanth gum, flash-frozen in liquid nitrogen-chilled isopentane and stored at –80°C. For all assays, samples were collected from matched regions of the same muscles by collecting cryosections (Leica CM1950 cryostat) for RNA analyses or histology and immunostaining assays.

### RNA extraction, RNA library preparation, RNA sequencing and bioinformatic analyses

Total RNA was extracted using TRIzol RNA isolation (Life Technologies) from frozen muscle samples. RNA was purified using RNeasy mini elute columns (Qiagen) and DNAse treated using Turbo DNA-free kit (Invitrogen). Purified, DNAse-treated RNA was quantified by NanoDrop and quality was assessed using Qbit RNA Assays (Thermofisher) and BioAnalyzer nano chips (Agilent) (RIN>7.8, Average 8.3±0.32). RNAseq library preparation and sequencing was performed using the TruSeq mRNA stranded kit (Illumina) and the Illumina HiSeq4000 Flow Cell with an average coverage of 63.05 million read pairs per sample at 2×75 base pair read length. The quality of the raw fastq reads from sequencer were evaluated using FastQC version 0.11.5 followed by adapter and quality trimming using Trimgalore. STAR 2.5.3a was used to map the reads to the reference mouse genome (GRCm38-mm10). The mapped reads were counted using HTSeq (version v0.11.0) with a reference genomic feature file (Gene transfer format, GTF). Overall analysis summary reports were analyzed using MultiQC v1.6. Differential gene expression analysis was performed using default parameters with Deseq2 version 1.26 (R 3.6). Visualization was performed by PCA, pheatmap, EnhancedVolcano and ggplot2 R packages. We set a threshold for log_2_ fold change (Log_2_FC) change of greater than an absolute value of 0.6 to select for the genes with significant differential expression. Gene lists were sorted by Log_2_FC (highest to lowest) to obtain the ranked list of differentially expressed genes and a p_adj_ value cutoff of 0.05 was used to assess statistical significance.

### Gene Set Enrichment Analysis (GSEA)

DESeq2 pairwise comparison results were filtered for 0.6 log_2_FC and 0.05 adjusted p_adj_ value and exported as tab delimited rank files (.rnk) files for upload into the stand-alone desktop version of GSEA (v4.1.0). The GSEA pre-ranked analysis was used with most default parameters except the following: the Ranked list = pairwise comparison “.rnk” file, Gene sets database = “c5.bp.v7.4.symbols” (Hallmark gene sets, GO biological processes, and gene symbols) and the Chip platform = “Mouse_Gene_Symbol_Re mapping_to_Human_Orth ologs_MSigDB.v7.4.chip”.

### Gene ontology analysis with Cytoscape and EnrichmentMap

The GSEA pairwise comparison results were uploaded into Cytoscape (v3.9.0) and analyzed using the EnrichmentMap pipeline collection plugins (v1.1.0). Comparisons were loaded into EnrichmentMap and the AutoAnnotate function was used with the MCL Cluster Annotation algorithm set to 5 words. The results are networks of related GO terms that are grouped together into a named network based on the most common works in each GO term within.

Autogenerated names of networks were renamed to fix grammar and nodes arranged to improve legibility. The leading-edge genes for each cluster (node) from the immune and extracellular matrix networks were exported for further analysis.

### Histology and histological analyses

Frozen muscles were removed from –80°C cryostorage and sectioned at an 8 μm thickness using a Leica CM1950 cryostat chilled to –20°C, where tissues were then mounted on slides and stained using Hematoxylin and Eosin (H&E), Alizarin Red, Picosirius red, and Masson’s Trichrome according to TREAT-NMD Standard Operating Procedures (SOPs) ^41^. Thresholding parameters were applied uniformly to whole cross-section tiled images acquired on the Olympus VS120-S5 Virtual Slide Scanning System using CellSens Version 1.13 and ImageJ FIJI Version 2.1./1.53c. For H&E-stained sections, areas of damage were selected using CellSens and quantified as percent damaged tissue area per total cross-sectional muscle area. For Alizarin Red stained sections, calcified areas were selected and quantified using CellSens as percent calcified tissue area per total cross-sectional muscle area. For Masson’s Trichrome stained sections, areas of fibrosis were calculated using FIJI (Image J) and reported as percent fibrosis per total cross-sectional muscle area ^41^.

### Immunofluorescence

Frozen muscles were removed from –80°C cryostorage and sectioned at an 8 μm thickness and mounted on slides for immunostaining procedures. Muscle sections were stained with anti-F4/80 (1:100, MCA497R, Bio-Rad), anti-COL1A1 (1:100, ab21286, Abcam), and anti-GAL-3 (1:100, ab76245, Abcam). First, muscle sections were fixed in ice-cold PFA for 10 min, washed in PBS (0.1% Tween-20), and blocked for 1 h in PBS supplemented with 10% goat serum (GeneTex), 0.1% Tween-20 (Sigma-Aldrich), and 10 mg/mL BSA (Sigma-Aldrich). Then sections were incubated with primary antibodies overnight at 4°C and subsequently probed with Alexa Fluor secondary antibodies, including goat anti-rat (H+L) Alexa Fluor 647 (1:500, A-21247, Thermo Fisher), goat anti-rabbit (H+L) Alexa Fluor 488 (1:500, A-11008, Thermo Fisher), and goat anti-rabbit (H+L) Alexa Fluor 568 (1:500, A-11011, Thermo Fisher). Sections were counterstained with wheat germ agglutinin (WGA) Alexafluor-647 (1:500, W32466, Thermo Fisher) to delineate cardiomyocytes and ProLong Gold Antifade with DAPI (P36935, Thermo Fisher) for nuclear staining.

### Gene expression analysis

Hearts from juvenile and adult dystrophic mice were used to perform gene expression analysis. In brief, total RNA was extracted from muscle samples by standard TRIzol (Life Technologies) isolation. Purified RNA (1000ng) was reverse-transcribed using Random Hexamers and High-Capacity cDNA Reverse Transcription Kit (Thermo Fisher Scientific). The mRNAs were then quantified using individual TaqMan assays on an ABI QuantStudio 7 Real-Time PCR machine (Applied Biosystems) using TaqMan Fast Advanced Master Mix (Thermo Fisher Scientific). Specific mRNA transcript levels were quantified using individual TaqMan assays (Thermo Fisher) specific for each mRNA target, including Ccl8 (Mm01297183_m1-FAM), Ccl3 (Mm00441259_g1_FAM-MGB), Ccl2 (Mm00441242_m1_FAM-MGB), Il1b (Mm00434228_m1_FAM-MGB), Anxa1 (Mm00440225_m1_FAM-MGB), Lgals3 (Mm00802901_m1_FAM-MGB), Stab2 (Mm00454684_m1_FAM-MGB), Arg1 (Mm00475988_m1_FAM-MGB), Il7r (Mm00434295_m1_FAM-MGB), Adam8 (Mm01163449_g1_FAM-MGB), Trem2 (Mm04209424_g1_FAM-MGB), Fpr2 (Mm00484464_s1_FAM-MGB), Spp1 (Mm00436767_m1_FAM-MGB), Fn1 (Mm01256744_m1_FAM-MGB), Col1a1 (Mm00801666_g1_FAM-MGB), Itgax (Mm00498701_m1_FAM-MGB), Mmp12 (Mm00500554_m1_FAM-MGB), and Timp1 (Mm01341361_m1_FAM-MGB). Gene expression for all mRNA targets was normalized to internal Hprt mRNA transcript levels using Hrpt Taqman assay (Mm03024075_m1_VIC-MGB).

### Pro-resolution preclinical drug trial

D2-*mdx* mice (18-19 days-old, n=6, males and females) were treated daily with pro-resolving, FPR2 agonist BMS-986235 (6mg/kg, oral gavage; HY-131180, MedChemExpress) for 3 weeks. Control D2-*mdx* mice were administered saline (18– 19-day-old, n=6, males and females) for 3 weeks. BMS-986235 was initially resuspended in 10% DMSO (Sigma) and 90% corn oil (Sigma), and further diluted in cherry syrup (NDC-0395-2662-16, Humco) for oral gavage. After treatment, hearts were harvested, imaged, flash-frozen in liquid nitrogen-chilled isopentane and stored at –80°C for molecular and histopathological analyses.

### Microscopy

We used Olympus VS120-S5 Virtual Slide Scanning System with UPlanSApo 40×/0.95 objective, Olympus XM10 monochrome camera, and Olympus VS-ASW FL 2.7 imaging software. Analysis was performed using Olympus CellSens 1.13 and ImageJ FIJI Version 2.1./1.53c software (National Institutes of Health). Brightfield whole tissue imaging was performed using Labomed Luxeo 6Z Digital HD Stereo Microscope with 10x/22mm objective and camera system.

**Statistics.** GraphPad Prism 9.2.0 was used for all statistical analyses of data. Statistical analysis was performed using non-parametric Mann-Whitney tests. Data normality was assessed for all statistical comparisons. All p-values less than 0.05 were considered statistically significant; *p < 0.05, **p < 0.01, ***p < 0.001, and ****p < 0.0001. Data plots reported as scatter plot with mean ± SD.

## Results

### D2-mdx model Exhibits Pediatric-Onset Cardiac Damage

We have previously described the use of D2-*mdx* model of DMD to define mechanisms contributing to pediatric-onset severe skeletal muscle degeneration ^41, 42^. Here we observed extensive pericardial damage of the ventricular wall in juvenile (<6-week-old) D2-*mdx* hearts (**Fig. 1A**). In contrast, the age-matched milder DMD model, B10-*mdx*, did not demonstrate any conspicuous histopathology (**Fig. 1B**). In D2-*mdx,* the histopathological damage extended to both the right and the left ventricular walls (**Fig. 1C**). Use of Sirius Red staining revealed extensive fibrosis, which manifests at notable levels in both the endomysium and perimysium of the ventricular walls in D2-*mdx* (**Fig. 1D**). Whole tissue cross-sectional analysis of H&E and Sirius Red stained hearts revealed increased cardiac wall damage and cardiomyocyte degeneration (*p<0.001*) (**Fig. 1C, E, F**), and a concomitant increase in fibrosis (*p<0.05*) in the juvenile D2-*mdx* hearts compared to age-matched B10-*mdx* hearts (**Fig. 1D, G, H**). The extent of damage in D2-*mdx* mice varied between the mice, reaching up to nearly 20% of total cardiac muscle cross-sectional area in the most affected case (**Fig. 1E, F**). Reminiscent of the skeletal muscle damage in D2-*mdx*, the cardiac muscle also exhibited increased calcification in damaged areas, which was exclusive to juvenile D2-*mdx* hearts (**Supplemental Fig. 1**). With mixed reports of cardiac pathology in older DBA2/J wildtype (D2-WT) hearts^43, 44^, we assessed if the juvenile-onset cardiac damage was also a feature of the juvenile D2-WT mice. Hearts from juvenile D2-WT showed no signs of gross pathology and were comparable to the hearts from the age-matched B10 wildtype (B10-WT) mice (**Supplemental Fig. 2A, B**). This was further confirmed by the microscopic evaluation that showed absence of any endomysial or perimysial fibrosis or calcification in the D2-WT hearts throughout the left and right ventricular walls (**Supplemental Fig. 2C, D**). Overall, these histopathological analyses reveal early onset spontaneous cardiac damage with significant fibro-calcification in the D2-*mdx*, offering a model to investigate the mechanisms of pediatric-onset cardiac damage and accompanying endomysial fibrosis observed in DMD.

**Figure 1.**
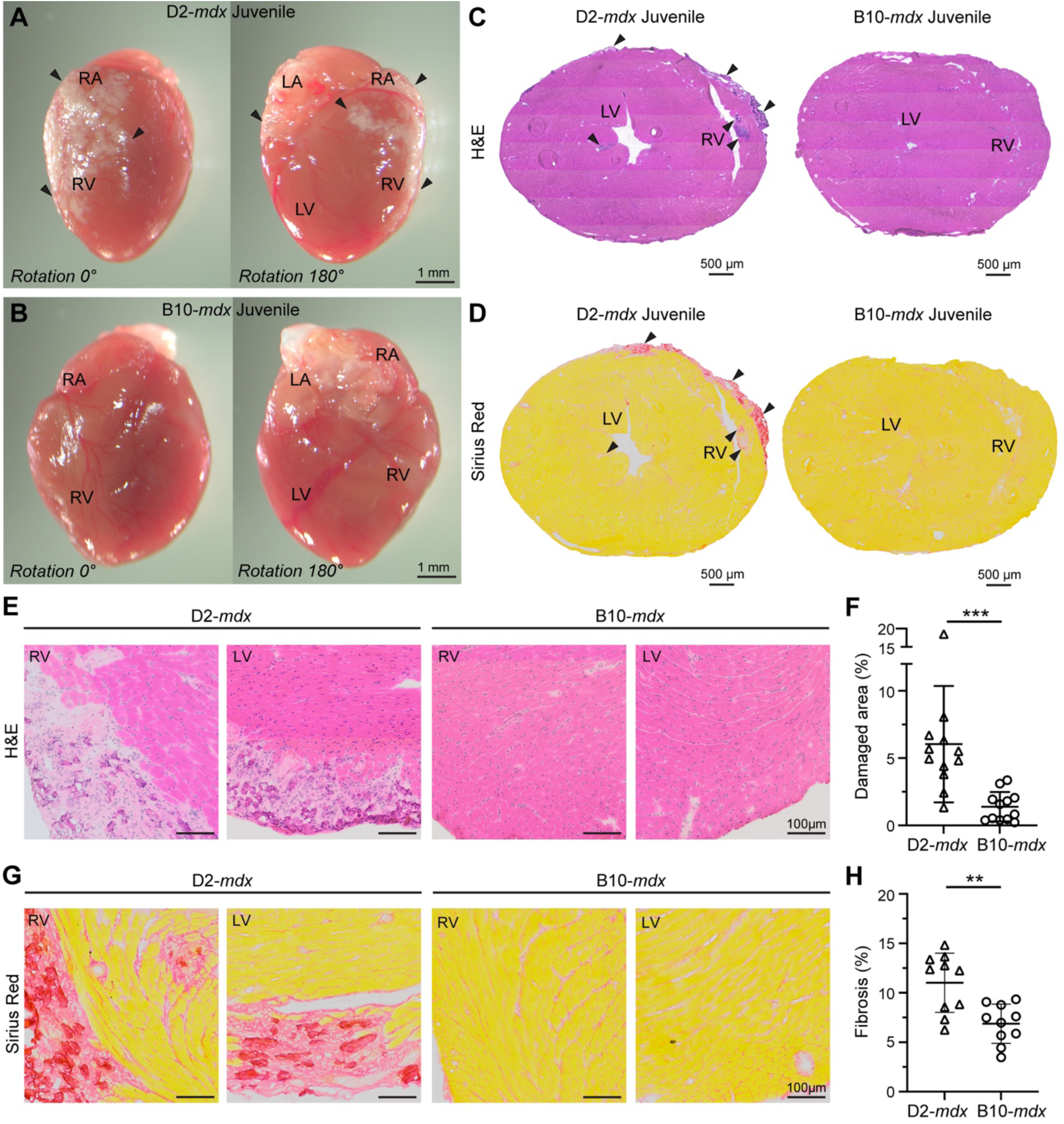
Histopathology associated with disease onset in juvenile D2-*mdx* hearts. Images show hearts harvested from juvenile (6 ± 0.5 wk) D2-*mdx* and B10-*mdx* mice at disease onset. **A-B.** Whole tissue images with matched orientation of D2-*mdx* (**A**) and B10-*mdx* (**B**) hearts showing ventricular and atrial fibro-calcified damage. **C-D.** Cross-sectional images of juvenile D2-*mdx* and B10-*mdx* hearts through the ventricular lumen, stained for histological features by H&E (**C**), and for fibrosis by Sirius Red (**D**). Arrowheads mark areas of fibrosis. **E-F.** Image (**E**) and quantification (**F**) showing a portion of heart cross-section from juvenile D2-*mdx* and B10-*mdx* hearts, showing damaged tissue areas characterized by the presence of interstitial fibrosis, inflammatory cells and damaged cardiomyocytes. **G-H.** Image (**G**) and quantification (**H**) showing a portion of heart cross-section from juvenile D2-*mdx* and B10-*mdx* hearts labeled with Sirius Red to mark fibrotic tissue area in hearts from juvenile D2-*mdx* and B10-*mdx* mice. Data represent mean ± SD from *n* = 12 hearts per cohort. Statistical analyses performed using Mann Whitney U test; \**p* < 0.05, \*\**p* < 0.001, \*\*\**p* < 0.001. Refer to Supplementary Fig. 1-2 for additional details.

### Dysregulated inflammatory response characterize cardiac damage in juvenile D2-*mdx*

To identify the molecular alterations associated with the pediatric-versus adult-onset cardiac damage caused by dystrophin deficit, we performed a comparative transcriptomic analysis of hearts from D2-*mdx* and B10-*mdx* mice. Using bulk RNA sequencing we examined genes that are differentially expressed (Log_2_FC 0.6 and p_adj_ value 0.05) between the 6-week-old male B10-*mdx* and D2-*mdx* hearts. This identified a total of 5,344 differentially expressed genes (DEGs) representing 22.3% of all protein-coding murine genes. Principal component analysis (PCA) identified that gene expression profiles of D2-*mdx* hearts were distinctly segregated from B10-*mdx* along both PC1 (72% variance) and PC2 (15% variance) axes (**Fig. 2A**). Intra– and inter sample variability of DEGs by heatmap analysis of the top 2,719 DEGs revealed consistent trends for both up– and down-regulated genes based on the genotype at disease onset, with 1,586 genes upregulated in D2-*mdx* and 1,133 genes upregulated in B10-*mdx* (**Fig. 2B**).

**Figure 2.**
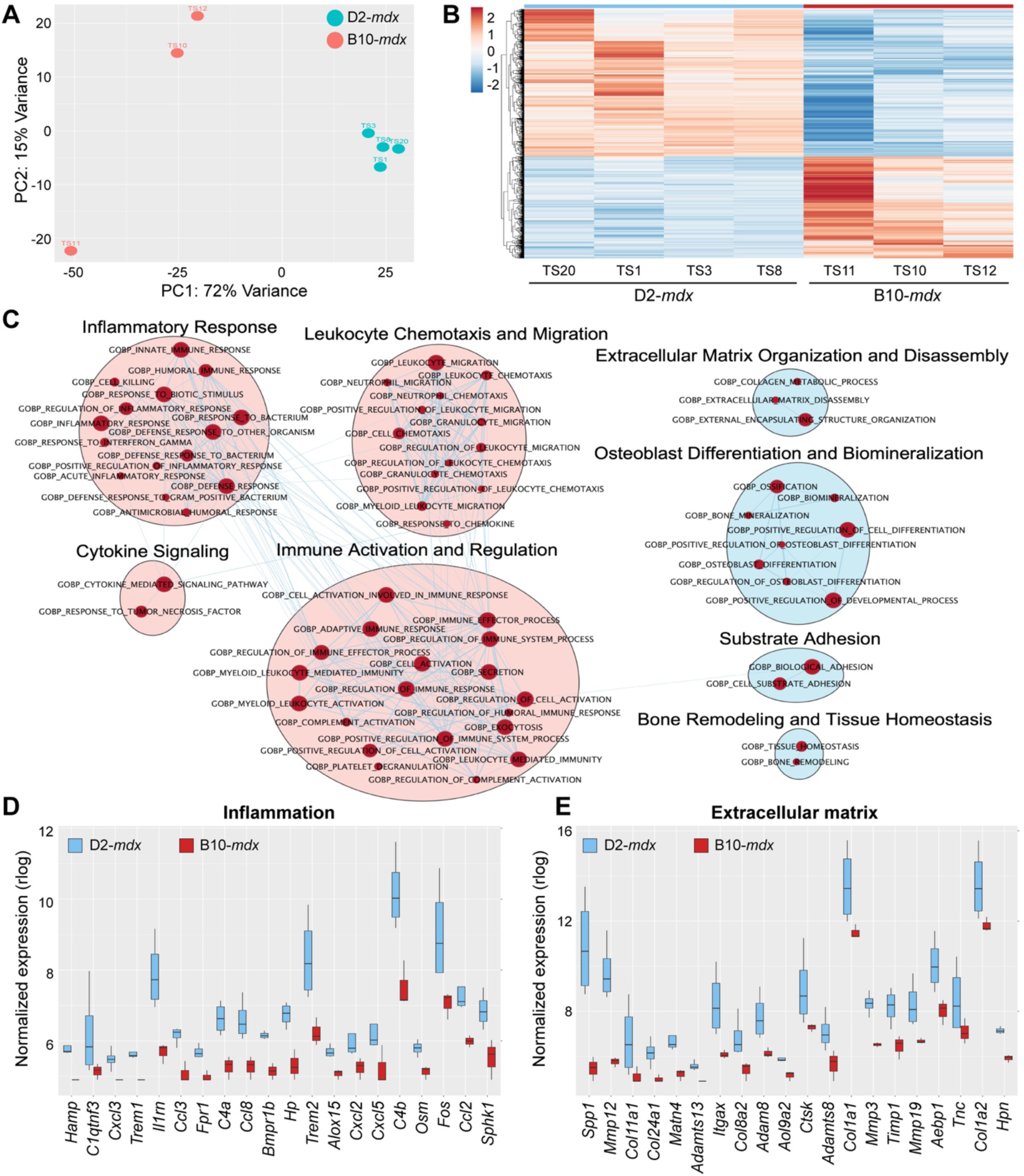
RNAseq analysis at disease onset in D2-*mdx* and age-matched B10-*mdx* hearts. Juvenile D2-*mdx* and B10-*mdx* hearts at disease onset (6 wk ± 0.5 wk) were analyzed by bulk tissue RNAseq. **A.** Dimensionality reduction of whole transcriptomic data via PCA (n=3-4 hearts/genotype) to assess sample clustering and inter-/intra-sample variance. **B.** Differential gene expression analysis depicted via heatmap plot of 2,719 DEGs observed between D2-*mdx* (blue) and B10-*mdx* (red), with 1,586 genes upregulated for D2-*mdx* and 1,133 genes upregulated for B10-*mdx*. Expression is z-score values of variance-stabilizing transformation (VST) normalized data. **C.** Gene Ontology (GO) analysis performed using Cytoscape and EnrichmentMap plugins to identify networks of related GO terms groups found upregulated (red dots) in juvenile D2-*mdx* hearts relative to B10-*mdx.* Pink clusters refer to inflammatory-related GO terms, while blue clusters refer to extracellular matrix-related GO terms. **D-E.** Boxplots showing VST normalized gene expression levels for top 20 differentially expressed inflammation-related (**D**; GOBP:Inflammatory Response) and extracellular matrix-related (**E**; GOBP:External Encapsulating Structure) transcripts observed between juvenile D2-*mdx* and B10-*mdx* hearts. Refer to Supplementary Fig. 3 for additional details.

To identify the functions of the DEGs and assess how they may contribute to the observed histopathology and functional deficit in juvenile D2-*mdx* hearts, we performed Gene Set Enrichment Analysis (GSEA) using our DESeq2 pairwise comparison results (0.6 log_2_FC and 0.05 adjusted p-value), followed by gene ontology (GO) analysis using Cytoscape’s EnrichmentMap pipeline (**Fig. 2C**). GO EnrichmentMap analysis predominantly indicated that upregulated DEGs in the juvenile D2-*mdx* fell within biological processes (BP) specific to immune response and extracellular matrix organization and remodeling. These included GOBP terms implicated in regulation of the inflammatory response, immune cell activation, leukocyte chemotaxis and migration, and cytokine signaling (**Fig. 2C**, red clusters), as well as extracellular matrix organization and disassembly, osteoblast differentiation, substrate adhesion and bone remodeling, and overall tissue homeostasis (**Fig. 2C**, blue clusters). Outputs from GSEA provided a comprehensive GO analysis of all upregulated and downregulated GOBP terms, their respective normalized enrichment scores (NES), p-values, and enrichment plots identifying alterations in inflammatory and extracellular matrix GOBPs have strongest positive enrichment scores (ES) (**Supplemental Table 1**, **Supplemental Fig. 3**). The top 20 upregulated GOBP terms identified dysregulation of the inflammatory response or extracellular matrix architecture that included a total of 331 upregulated DEGs common between them (**Supplemental Table 2**). This was validated by the quantification of the normalized expression for the top 20 most upregulated DEGs specific to unique GOBPs such as inflammatory response (GO:0006954) (**Fig. 2D**) and external encapsulating structure (extracellular matrix) organization (GO:0045229) (**Fig. 2E**). The leading-edge genes identified increased expression of pro-inflammatory chemokines, and components of fibrotic extracellular matrix remodeling as potential drivers of overt pediatric-onset cardiac damage in the D2-*mdx* (**Figure 2, Supplementary** Fig. 3).

To independently validate the role of aberrant acute inflammatory response of granulocyte and leukocyte chemotaxis by way of chemokine/cytokine signaling pathways, we examined these genes by qPCR in an expanded (n = 9 D2-*mdx* and n = 7 B10-*mdx*) cohort of juvenile *mdx* mice. This validated our findings from RNAseq analysis and confirmed significant upregulation of macrophage-secreted pro-inflammatory C-C family chemokines, *Ccl3* (macrophage inflammatory protein-1α) and *Ccl8* (monocyte chemoattractant protein-2), that regulate neutrophil, monocyte and lymphocyte chemotaxis following acute tissue damage (**Fig. 3A**; *p<0.001*). Similarly, expression of *Il7r* (interleukin-7 receptor), that promotes neutrophil and monocyte recruitment, was also found upregulated in D2-*mdx* hearts (**Fig. 3A**; *p<0.001*), while, expression of *Stab2* (stabilin-2), a macrophage-expressed phosphatidylserine (PS) surface receptor that mediates phagocytosis and extracellular matrix remodeling during inflammation ^45^, was also upregulated in D2-*mdx* hearts (**Fig. 3A**; *p<0.001*). Expression of *Adam8* (a disintegrin and metalloproteinase 8), that promotes release of pro-inflammatory cytokines and cell adhesion molecules and degradation of extracellular matrix ^46^, was also highly overexpressed in D2-*mdx* hearts in accordance with bulk RNAseq results (**Fig. 3A**; *p<0.001*). *Trem2* (triggering receptor expressed on myeloid cells 2), which drives NF-kB signaling and production of pro-inflammatory cytokines including IL-6 and TNFα ^47^, was also upregulated in juvenile D2-*mdx* hearts (**Fig. 3A**; *p<0.01*).

**Figure 3.**
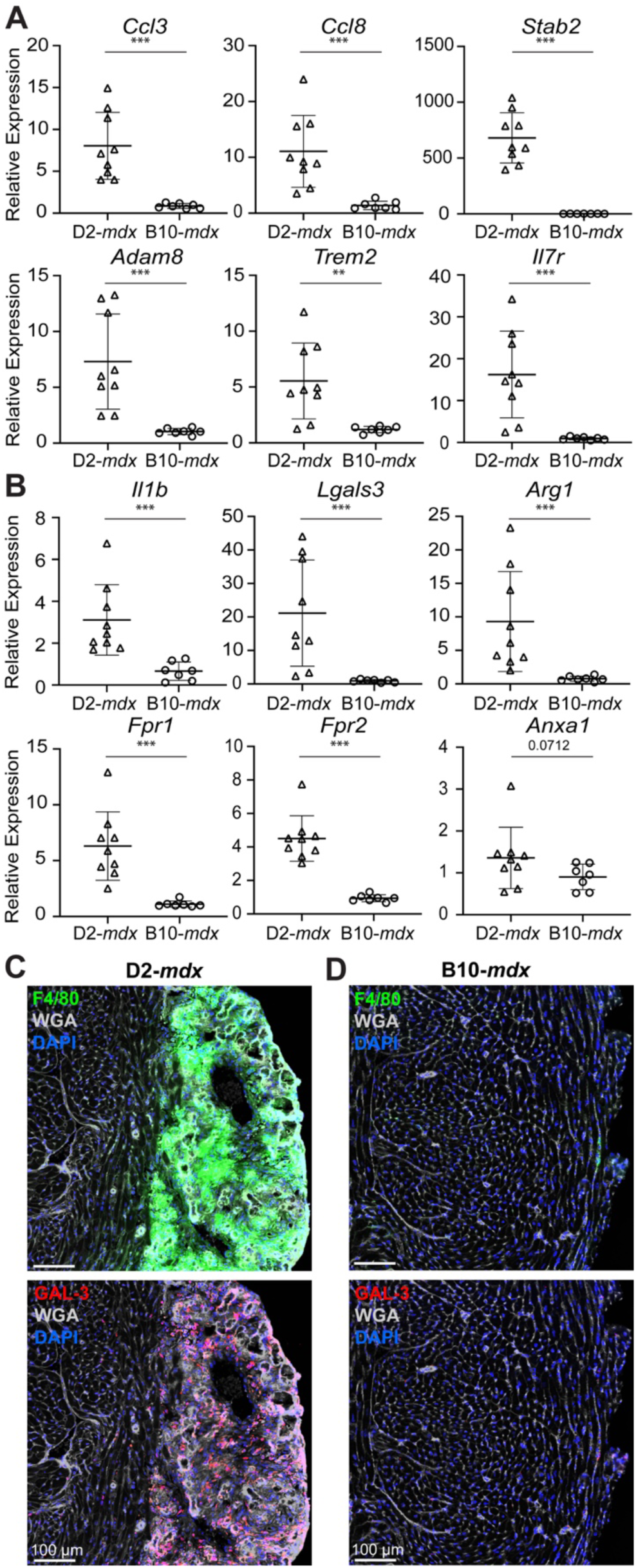
Targeted analysis of inflammatory response at disease onset in D2-*mdx* B10-*mdx* hearts. **A.** qRT-PCR analysis of a distinct cohort of D2-*mdx* and B10-*mdx* hearts to assess the expression of inflammatory genes identified by RNAseq analysis cohort to be differentially expressed. Transcripts include top dysregulated genes involved in leukocyte activation, migration and chemotaxis and regulation of inflammatory response and leukocyte-mediated immunity (*Ccl3, Ccl8, Stab2, Adam8, Trem2, Il7r*). Relative gene expression values normalized to internal *Hprt* transcript levels. **B.** qRT-PCR analysis of a distinct cohort of D2-*mdx* and B10-*mdx* hearts to assess inflammatory genes that show broad dysregulation of neutrophil and macrophage response in juvenile D2-*mdx* hearts. Relative gene expression values normalized to internal *Hprt* transcript levels. **C-D.** Images showing immunostaining for pan-macrophage marker, F4/80 (green), and pro-inflammatory, pathogenic macrophage marker, GAL-3 (red), in juvenile D2-*mdx* (**C**) and B10-*mdx* (**D**). Data represent mean ± SD from *n* = 7-9 hearts per cohort. Statistical analyses performed using Mann Whitney U test; \**p* < 0.05, \*\**p* < 0.001, \*\*\**p* < 0.001. For age-matched WT controls, refer to Supplementary Fig. 4.

With the abundant increase in inflammatory cell chemokines, we examined the abundance of macrophages using the pan-macrophage marker F4/80. This confirmed extensive presence of macrophages in the damaged regions in D2-*mdx* hearts, which is distinct from what is observed in the B10-*mdx* hearts (**Fig. 3C, D**; *top panels*). To assess the attributes of infiltrating macrophages in the D2-*mdx* hearts, we next profiled transcript levels of *Il-1b* (interleukin-1b), a pro-inflammatory macrophage marker, *Arg1* (arginase-1), a pro-regenerative macrophage marker, and *Lgals3* (galectin-3), a pathogenic macrophage marker^48, 49^. While the inflammatory and regenerative marker genes were increased by 3-, and 9-folds respectively in D2-*mdx*, compared to B10-*mdx* hearts, the levels of pathogenic marker (*Lgals3*) was elevated by ∼45-fold (*p<0.001*) (**Fig. 3B**). To determine if the different genetic background may contribute to this dysregulation we examined these transcripts in age-matched juvenile D2-WT and B10-WT mice. This showed no influence of genetic background for *Adam8*, *Il1b*, *Lgals3*, and *Arg1* between strains, and minimal impact on the expression of *Ccl3* (*p<0.01*), *Ccl8* (*p<0.01*), *Trem2* (*p<0.01*), and *Il7r* (*p<0.05*) (**Supplemental Fig. 4A**). As our previous investigations have revealed galectin-3 enriched macrophages as a driver of fibrotic degeneration of D2-*mdx* skeletal muscles^50^, we assessed the tissue localization and abundance galectin-3 protein (GAL-3) in D2-*mdx*. This identified that GAL-3^+^ macrophages were highly abundant and nearly exclusively localized within the damaged regions of D2-*mdx* hearts (**Fig. 3C**), while in accordance with qPCR results, these pathogenic macrophages were absent in B10-*mdx* hearts (**Fig. 3B**, **D**).

Increased expression of chemokines observed in these tissues explains the excess of macrophages in D2-*mdx*. However, during acute injury, the inflammatory response is controlled by a pro-resolving response that follows cytokine-mediated activation of inflammation and involves activation of the G protein coupled receptors including Formyl peptide receptors (FPRs). FPRs are activated by the endogenous ligands produced in response to tissue damage, including lipids (Resolvin D1, Lipoxin A4) and protein (Annexin A1; AnxA1) that bind FPR1/2^51, 52, 53^ and serve as the master switch at the site of damage that helps resolve the inflammation.

They do so by promoting macrophage skewing from pro-to anti-inflammatory fates and regulating signaling pathways that help clear immune cell infiltration by activating their apoptosis and non-phlogistic clearance as well ^53, 54, 55, 56, 57^. To assess if the excessive inflammatory responses in D2-*mdx* hearts is due to reduced FPR signaling, we examined the expression of FPRs (*Fpr1*, *Fpr2*) and its ligand *Anxa1*. This revealed over 4-fold upregulation of FPRs (both *Fpr1* and *Fpr2*) expression in D2-*mdx* relative to B10-*mdx* heart, but no change in the expression of *Anxa1* between D2-*mdx* and B10-*mdx* (**Fig. 3B**). These results validated the findings from the previous cohort used for bulk RNAseq analysis (**Fig. 2, Supplementary Table 1**). Together, they suggest reduced activation of FPR signaling hinders resolution of inflammation and may be a driver of the excessive inflammation seen in juvenile D2-*mdx* hearts, which in turn is linked to their fibro-degenerative state.

### Fibrotic ECM remodeling drives early onset cardiac fibrosis in juvenile D2-*mdx*

To evaluate the prominent upregulation of ECM remodeling pathways implicated by the RNAseq analysis, we performed qPCR for multiple extracellular matrix-associated components and remodeling enzymes identified by the analysis of our bulk RNAseq cohort. This validated the observed upregulation of *Fn1 (*fibronectin; *p<0.01), Col1a1 (*collagen 1A; *p=0.0712), and Itgax* (integrin alpha X; *p<0.001)* in D2-*mdx* hearts, relative to B10-*mdx* (**Fig. 4A**). Assessment of *Spp1* (osteopontin), a DMD genetic modifier that links inflammation to extracellular matrix assembly and fibrosis ^58, 59, 60, 61, 62^ and macrophage-expressed matrix remodeling enzyme *Mmp12 (*matrix metalloproteinase 12), both showed over 200-fold upregulation in D2-*mdx* hearts, compared to B10-*mdx* (**Fig. 4A**). While other ECM regulator and structural components including *Timp1 (*TIMP metallopeptidase inhibitor 1), *Itgax*, *Fn1*, and *Col1a1* showed between 4-40-fold upregulation in D2-*mdx* hearts (**Fig. 4A**). These validate the findings from the bulk RNAseq cohort and implicate a nexus of inflammatory-ECM dysregulation in pediatric-onset cardiac pathogenesis in the D2-*mdx*. To monitor the sites of fibrosis in juvenile D2-*mdx* hearts, we immunostained heart cross-sections for COL1A1 and co-stained with wheat germ agglutinin (WGA), which showed dense COL1A1 expression in D2-*mdx* in the damaged regions along the RV wall and throughout the endomysium (**Fig. 4B, B’**), while endomysial COL1A1 staining in B10-*mdx* counterparts was minimal in comparison (**Fig. 4C, C’**). To address the influence of genetic background on the above findings, we assessed expression in D2-WT and B10-WT hearts, which indicated no genotype-related differences in the expression of *Fn1*, *Col1a1*, *Itgax*, or *Timp1*, and a comparatively modest increases in the expression of *Spp1* (*p<0.01*) and *Mmp12* (*p<0.01*) in D2-WT hearts, relative to B10-WT (**Supplemental Fig. 4B**), when compared to differences between mdx strains (**Fig. 4A**). This identified the site and composition of the fibrotic ECM in the D2-*mdx* heart. It also highlighted the potential of targeting the aberrant pro-inflammatory response to attenuate fibrotic cardiac degeneration of the D2-*mdx* dystrophic heart.

**Figure 4.**
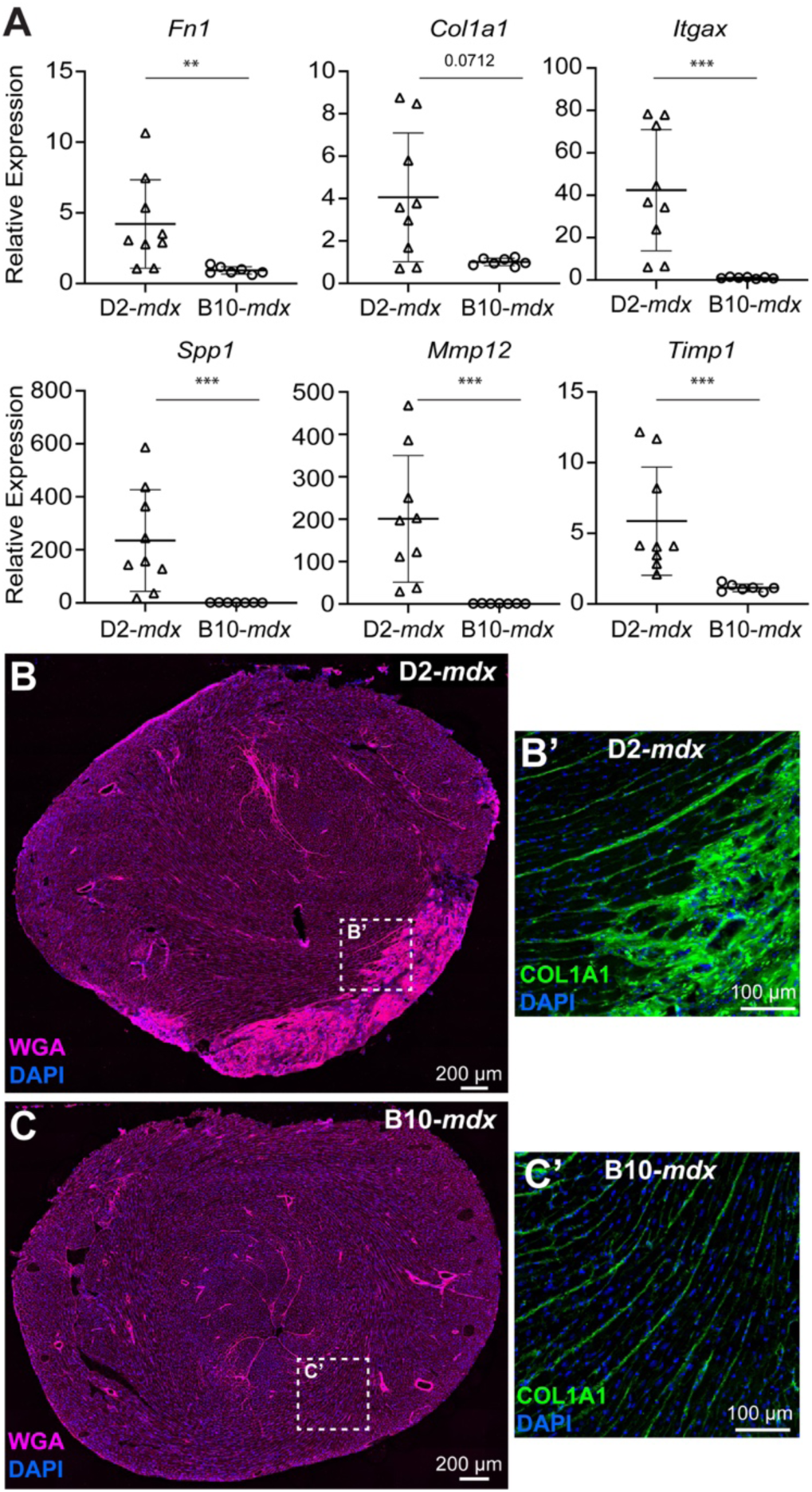
Targeted analysis of extracellular matrix remodeling response at disease onset in D2-*mdx* B10-*mdx* hearts. **A.** qRT-PCR analysis of a distinct cohort of D2-*mdx* and B10-*mdx* hearts to assess the expression of extracellular matrix-associated genes involved in matrix organization/re-organization (*Fn1, Col1a1, Itgax, Spp1, Mmp12, Timp1)* that are identified to be dysregulated by the RNAseq cohort. Relative gene expression values normalized to internal *Hprt* transcript levels. **B-C.** Images showing extracellular matrix distribution visualized using wheat germ agglutinin (WGA, pink) within, and surrounding, areas of cardiac damage in juvenile D2-*mdx* (**B**), and B10-*mdx* (**C**). **B’-C’.** Zoom of the dotted area from whole cross-sectional images showing immunostaining for COL1A1 within the extracellular matrix shows increased COL1A1 expression in damaged D2-*mdx* hearts (**B’**), relative to B10-*mdx* (**C’**), indicative of early-onset endomysial fibrosis. Data represent mean ± SD from *n* = 7-9 hearts per cohort. Statistical analyses performed using Mann Whitney U test; \**p* < 0.05, \*\**p* < 0.001, \*\*\**p* < 0.001. For age-matched WT controls, refer to Supplementary Fig. 4.

### Activation of pro-resolving FPR signaling prevents cardiac damage in D2-*mdx* hearts

The above findings link inflammation and cardiac fibrosis in D2-*mdx*. With mixed success of corticosteroid use in treating this in heart by dampening inflammation, coupled with our findings of poor activation of pro-resolving FPR signaling, we hypothesized use of FPR-targeting therapy may resolve the chronic inflammation, without blocking acute inflammation required for the reparative ability of the injured tissue^63, 64^ (**Fig. 5A**).

**Figure 5.**
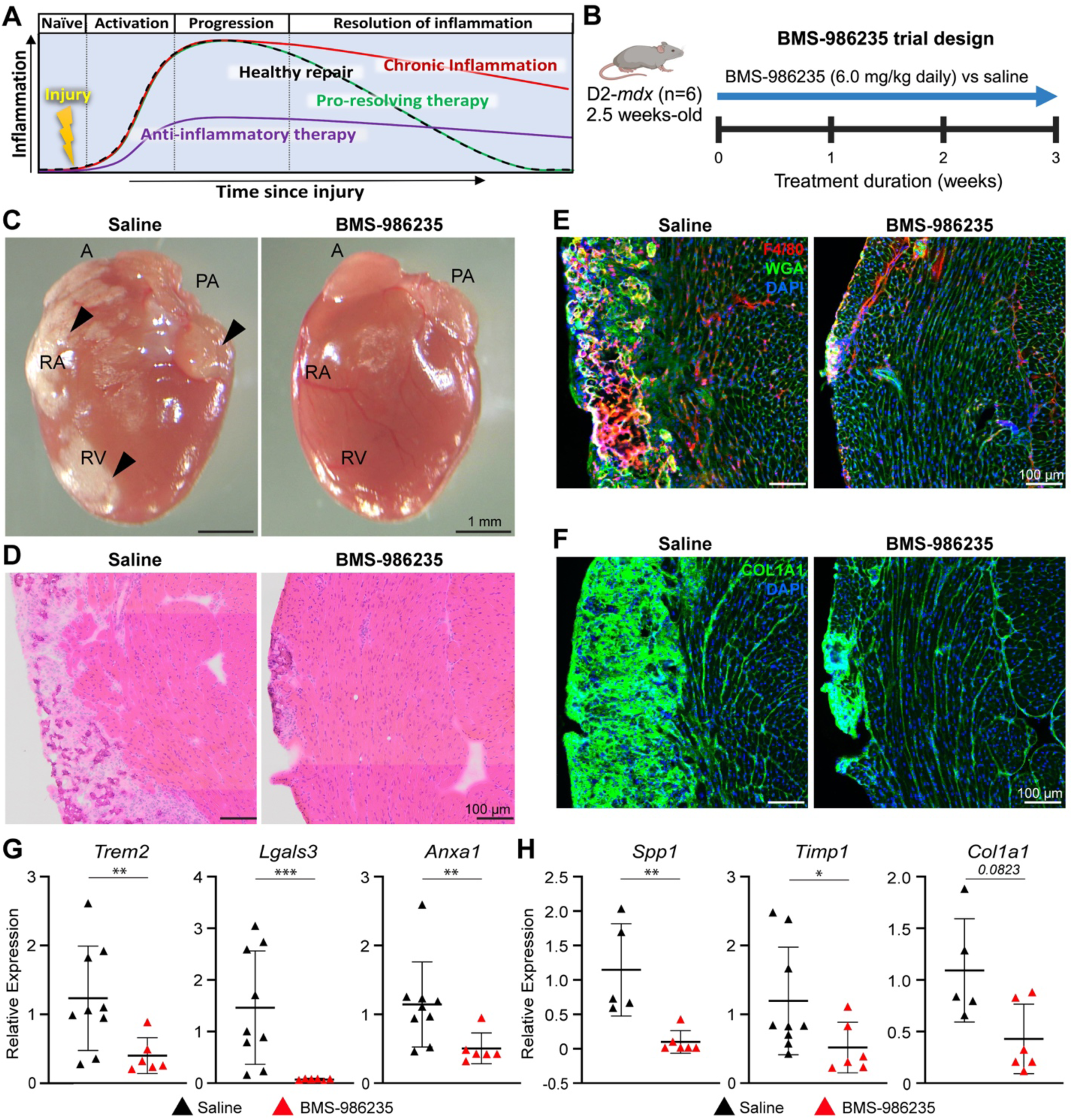
Pro-resolving therapy to mitigate cardiac disease onset in juvenile D2-*mdx*. **A.** Schematic describing the inflammatory response following cardiac injury in health (black trace) or dystrophic (red trace), showing acute versus chronic inflammation respectively. Use of anti-inflammatory drug (purple trace) lowers inflammatory response blunting inflammation, instead use of pro-resolving therapy (green trace) does not impact the onset of inflammation but helps clear inflammation preventing the inflammation to become chronic. **B.** Schematic detailing the pre-clinical testing of pro-resolving drug, BMS-986235 (6.0mg/kg, 3 wk daily administration) in D2-*mdx* mice (n=6) just prior to disease onset (18-19 days old). **C.** Whole tissue images of the matched orientation of hearts showing ventricular and atrial fibro-calcified damage in saline or BMS-986235-treated D2-*mdx* mice. **D.** Image showing cross-section of D2-*mdx* heart stained for histological features by H&E from mice treated with saline or BMS-986235. **E.** Images showing cross-section of D2-*mdx* heart immunostained for pan-macrophage marker, F4/80 (red) and counterstained with WGA (green) and DAPI (blue) to mark the ECM and nuclei, respectively. **F.** Image showing heart cross-section immunostained for COL1A1 and counterstained with DAPI (blue) to visualize nuclei. **G-H.** qRT-PCR analysis of inflammatory (**G**) and extracellular matrix genes (**H**) to assess the effect of drug treatment of D2-*mdx* mice (red triangles), as compared with saline-treated controls (black triangles). Relative gene expression values normalized to internal *Hprt* transcript levels. Data represent mean ± SD from *n* = 6 hearts per cohort. Statistical analyses performed using Mann Whitney U test; \**p* < 0.05, \*\**p* < 0.001, \*\*\**p* < 0.001.

To assess the benefit of a pro-resolving FPR-agonist therapy for pediatric-onset cardiac pathology in D2-*mdx*, we tested use of the synthetic FPR agonist BMS-986235 versus saline (n = 6 animals/cohort). Mice were orally dosed with drug beginning at 2.5-weeks of age (just prior to the onset of cardiac pathology) and were maintained on drug (or saline) till 6-weeks of age when tissues were harvested for further analysis (**Fig. 5B**). Analysis of histopathology of the cardiac tissue cross-section revealed a clear lack of fibro-calcified damaged areas along the RV and LV walls in drug-treated hearts, compared to saline controls (**Fig. 5C**). Histological analyses performed by H&E staining, confirmed this observed therapeutic effect with some of the drug-treated hearts showing only small and discrete areas of damage within the RV wall, and the rest lacked any signs of damage or inflammation altogether, while the saline-treated mice showed extensive cardiac damage (**Fig. 5D**). To further assess this impact of pro-resolving therapy on inflammation and extracellular matrix remodeling in D2-*mdx* hearts, we immunostained tissue cross-sections for macrophages, which confirmed the reduction in macrophage infiltration through the heart and within and surrounding any small sites of damage that existed in our treated cohort (**Fig. 5E**). This was in stark contrast to the heightened macrophage infiltration both within and surrounding sites of damage in saline control hearts (**Fig. 5E**). Next, to more directly assess fibrotic gene expression we immunostained these hearts for COL1A1 and found the drug treatment also significantly reduced the COL1A1 expression throughout the heart, when compared to the expression in the saline controls (**Fig. 5F**).

To assess the effect of acute BMS-986235 treatment on inflammation and extracellular matrix remodeling pathways, we next performed targeted qPCR for both inflammatory– and extracellular matrix remodeling-associated transcripts previously shown to be dysregulated in D2-*mdx* hearts (**Fig. 2-4**). We found acute BMS-986235 treatment of juvenile D2-*mdx* mice resulted in a significant reduction in the levels of inflammatory transcripts *Trem2*, *Lgals3*, and *Anxa1* (**Fig. 5G**), confirming the efficacy of this pro-resolving therapy to attenuate aberrant inflammatory signaling via the FPR2-ANXA1 axis. Similarly, quantification of extracellular matrix remodeling targets, *Spp1*, *Timp1,* and *Col1a1*, showed significant depletion of *Spp1* and *Timp1* transcripts (**Fig. 5H**), and a trending reduction in *Col1a1* transcript levels (**Fig. 5H**) which aligns with COL1A1 immunostaining results (**Fig. 5F**). Future chronic studies will be required to assess the full benefit of chronic pro-resolving therapy to delay the onset and lessen the severity of fibrotic cardiac degeneration with disease progression in older D2-*mdx* mice.

## Discussion

While there has been a long-standing recognition of early-onset cardiac deficit and its lethal consequences for boys with DMD, our experimental understanding of the molecular deficits and preclinical interventions have been based on the use of adult mouse models. This is due to the late onset of cardiac damage in the *mdx* mouse model. Our studies here introduce D2-*mdx* as a model that manifests pediatric-onset cardiac damage. Histopathological analyses reveal early onset spontaneous cardiac damage with significant fibro-calcification in the D2-*mdx* model, offering an opportunity to investigate the mechanisms of pediatric-onset cardiac damage and accompanying endomysial fibrosis observed in DMD.

Our studies here identify dysregulation of inflammatory and ECM remodeling as two such pathological pathways. Next, we focused on harnessing underlying molecular pathways that distinguish the cardiac deficit in D2-*mdx* model as compared to the adult-onset cardiac deficit B10-*mdx* model. This analysis allowed distinguishing the differences driving the onset of mild and severe cardiac muscle degeneration and cardiomyopathy independent of the presence/absence of dystrophin protein. Such differences are reminiscent of the DMD patients, who manifest varying level and severity of cardiomyopathy despite the lacking dystrophin expression.

Our studies identify excessive inflammatory response that fails to resolve through FPR2-mediated pro-resolving pathway prevents restoring the injured heart tissues to their uninflamed state. While the infiltrating leukocytes are needed in damaged heart to clear the dead cells, FPR2 and other mediators that repress inflammation are released leading to predominance of anti-inflammatory cells – a response associated with activation of cardiac repair. We find that this latter repressive response is poor, and that the excessive inflammatory signaling proceeds via Spp1 and TGFβ pathway to activate downstream cardiac fibrosis and other degenerative response. Activation of these profibrotic degenerative response have long been recognized as a feature of the D2-*mdx* model^65^ which we previously showed contributes to excessive skeletal muscle pathology in the juvenile D2-*mdx* mice by suppression of skeletal muscle regeneration^41,42^. Unlike skeletal muscle, cardiac muscle does not undergo regeneration and our comparative analysis of D2-*mdx* and B10-*mdx* identify cardiomyocyte degeneration due to chronic inflammation and loss by way of fibro-calcified ECM. We observe the ventricular pericardium as the region most affected by this damage, but this can progress to the atria as well (**Fig. 1**). This variability in the affected region is in addition to the variability we observe in the severity of cardiac damage between individual mice. This hints at a level of stochasticity in the degeneration. We believe this may plausibly reflect the level of initial damage to the affected heart, which is then amplified as the inflammation ensues becoming chronic.

We find chronic cardiac inflammation in D2-*mdx* is marked by higher level of activation of inflammatory and immune response in part by higher expression of chemokines that attract these immune cells (**Fig. 2, 3**). This inflammatory response involves accumulation of *Spp1*(OPN)/*Lgals3*(GAL-3) expressing pathogenic macrophages that we recently identified by single cell RNAseq analysis of the skeletal muscles from *mdx* mice^49^. Osteopontin secreted by these macrophages promote skeletal muscle fibrosis by activating the stromal progenitors and we suggest a similar mechanism may be in place following accumulation of these macrophage in the damaged heart (**Fig. 3**). In support of this mechanism, we find these GAL-3^+^ macrophages enriched in the same pericardial region that are enriched in COLA1 indicative of active fibrosis (**Fig. 3**). The indication that this is an active phenomenon comes from the concomitant enrichment of other ECM building (fibronectin) and degrading (TIMP/MMP) components along with leukocyte attracting (CCCL3/8) and resolution (FPR1/2) signaling (**Fig. 2-4**). This dynamic indicated that there is a likely shift in the equilibrium of these two opposing – inflammation building and resolving signals, and hence rebalancing this could provide a likely beneficial effect for the affected region. This is in line with the fact that while acute inflammation following tissue damage is essential for cardiac repair, chronic inflammation can be disruptive^66^. To address this ensuing imbalance of these two opposing inflammatory signaling we made use of the pro-resolving therapy that unlike GCs, precisely targets the resolution of inflammation by activating FPR2 signaling, but not blocking activation of inflammation by NFkb or related pro-inflammatory pathways. This approach showed excellent promise such that a short (3 week) treatment of juvenile D2-*mdx* mice allowed full resolution of inflammation and prevented any subsequent fibrotic cardiac damage detected histologically as well as by way of aberrant molecular signature including *Gal3*/*Spp1* macrophages and *Col1A1* and *Fn1* expressing stromal cells (**Fig. 5**).

In summary, our studies introduce the D2-*mdx* as a model that manifests pediatric-onset cardiac damage, which provides opportunities to investigate the drivers of early-onset cardiomyopathy in DMD patients. We identified dysregulation of inflammatory and ECM remodeling pathways as key contributors. Specifically, our findings highlight excessive and unresolved inflammatory response involving pathogenic macrophages and neutrophils, contributes to chronic inflammation and progressive cardiomyocyte degeneration and fibrotic replacement in the D2-*mdx* model. Finally, our use of FPR2-targeting therapy provides a potential therapeutic avenue to prevent or mitigate pediatric-onset cardiac pathologies in DMD.

## Author Contributions

This study was conceived, and the experiments designed by JSN and JKJ with inputs from all authors. RNAseq sample processing was conducted by JSN, RH, DAM, and KP; bioinformatic analyses were performed by AL, PU, SB, ST, and VB. AL conducted molecular analyses with help from YJM and ST. Histopathological analyses were conducted by AL, RH, YJM, IHG, and GW. The manuscript was written by JSN, AL, and JKJ, and edited by all authors. JKJ and JSN obtained funding and provided oversight for pursuit of the study.

## Acknowledgements

This work was supported by funds provided by Children’s National Research Institute (JSN) and the Foundation to Eradicate Duchenne (JKJ; JSN). JKJ and JSN acknowledge support from the Department of Defense DMDRP (W81XWH2110711| JSN; W81XWH2110680 | JKJ) for their work on DMD. Microscopy was performed at the Cell and Tissue Microscopy Core supported by CNRI and The National Institutes of Health NICHD (P50HD105328 | JKJ).

## Supplemental Material

**Supplemental Figure 1.**
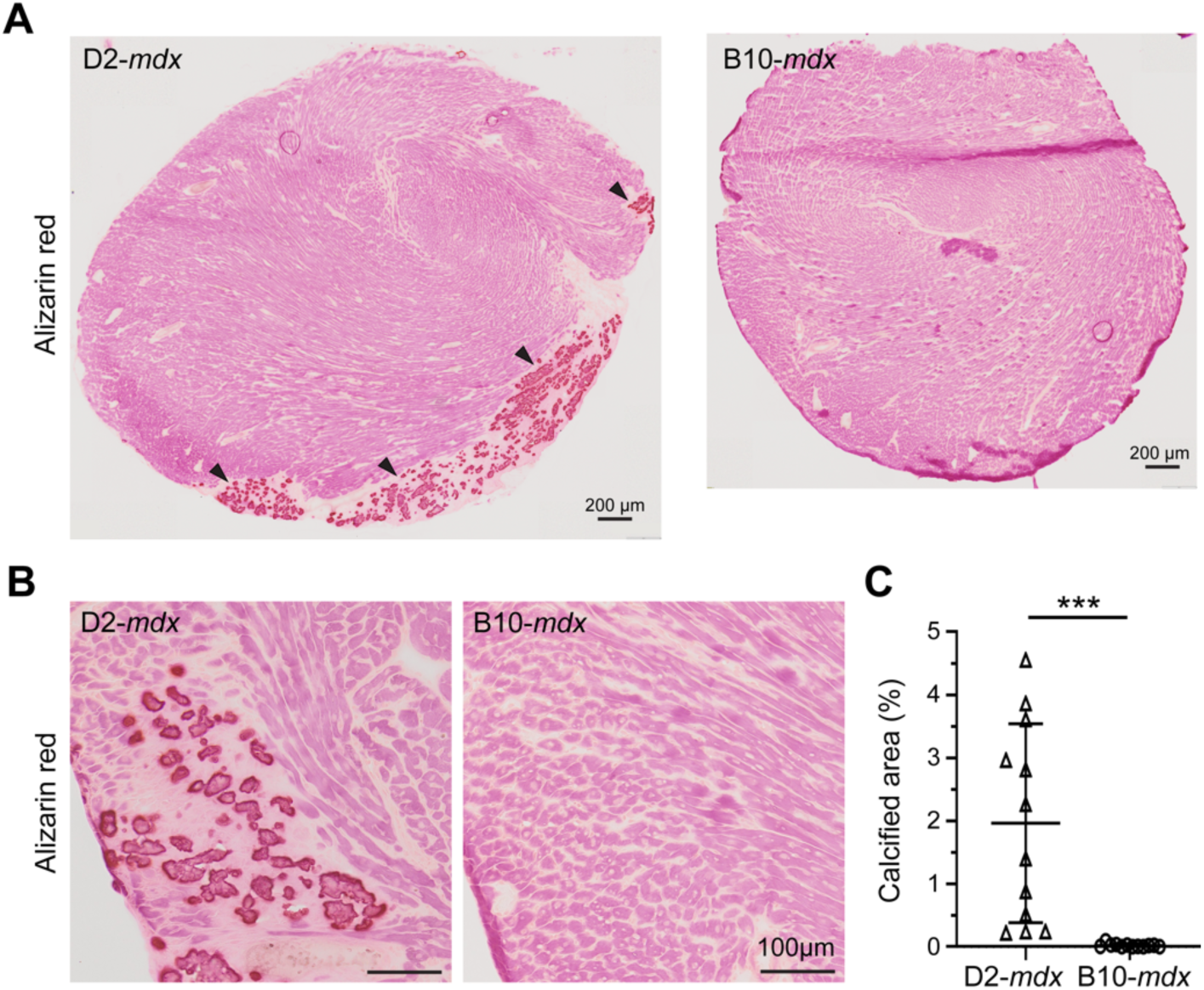
Cardiac histopathology in juvenile D2-*mdx* model. **A.** Alizarin Red staining of juvenile D2-*mdx* and B10-*mdx* hearts (whole cross-section) showing right ventricular (RV) heart damage and calcification in juvenile D2-*mdx* hearts. Scale bars indicate 200µm. **B-C.** High magnification images from panel A, showing Alizarin Red staining of juvenile D2-*mdx* and B10-*mdx* hearts, and corresponding quantification of calcified fiber area per total tissue area. of fibrosis, damage and calcification are highlighted by black arrowheads (**C**). Scale bars indicate 100µm. Statistical analyses performed using Mann Whitney U test; \*\*\**p* < 0.001.

**Supplemental Figure 2.**
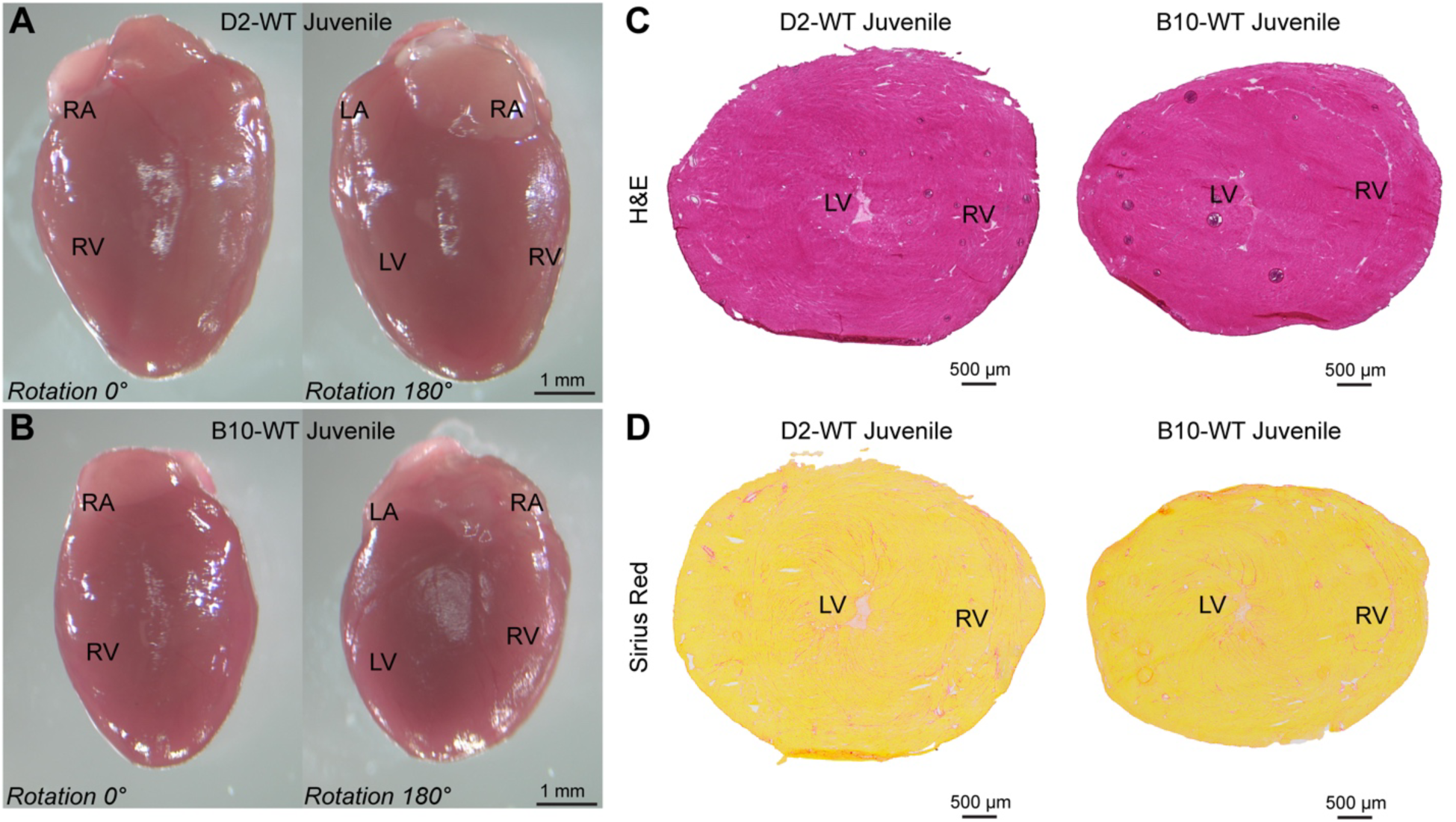
Histopathological assessment of juvenile D2-WT and B10-WT hearts. Images show hearts harvested from juvenile (6 ± 0.5 wk) D2-WT and B10-WT mice. **A-B.** Whole tissue images with matched orientation of D2-WT (**A**) and B10-WT (**B**) hearts showing lack of any ventricular and atrial fibro-calcified damage. **C-D.** Cross-sectional images of juvenile D2-*mdx* and B10-*mdx* hearts through the ventricular lumen, stained for histological features by H&E (**C**), and for fibrosis by Sirius Red (**D**). Both D2-WT and B10-WT hearts lack any signs of fibro-calcified damage.

**Supplemental Figure 3.**
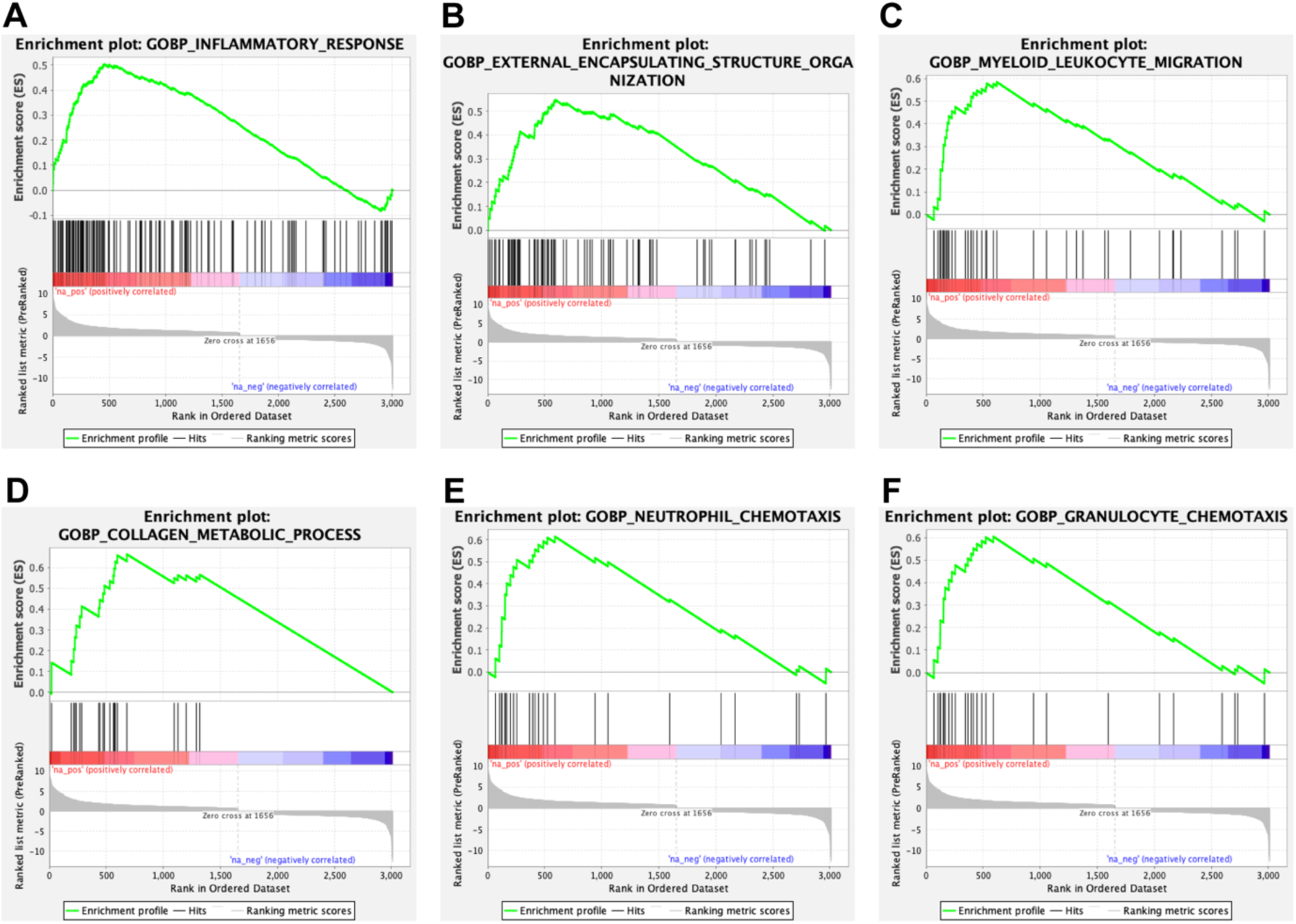
GSEA enrichment plot analysis of top gene ontology (GO) hits from differential gene expression analysis in juvenile D2-*mdx* and B10-*mdx* hearts. **A-F.** Enrichment plot analysis of top immune-related (**A, C, E, F**) and extracellular matrix-related (**B, D**) GO biological process (BP) terms obtained using GSEA, including Inflammatory Response (**A**), Encapsulating Structure Organization (**B**), Myeloid Luekocyte Migration (**C**), Collagen Metabolic Process (**D**), Neutrophil Chemotaxis (**E**), and Granulocyte Chemotaxis (**F**).

**Supplemental Figure 4.**
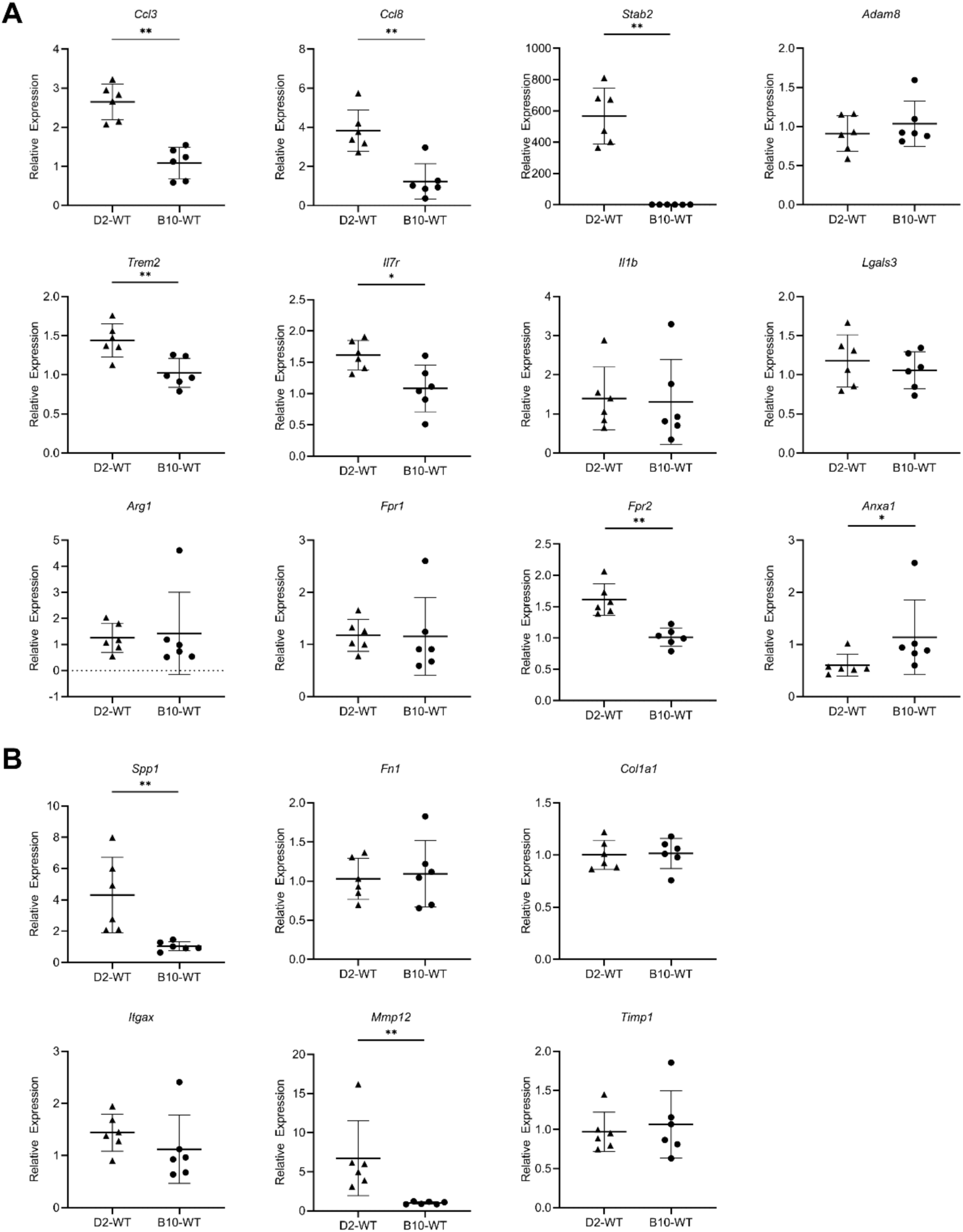
Gene expression analysis of D2-WT and B10-WT hearts for selected immune and extracellular matrix targets. **A.** Relative gene expression analysis for dysregulated immune-related genes between D2-*mdx* vs. B10-*mdx* (**Figure 3**) assessed in juvenile D2-WT and B10-WT hearts. **B.** Relative gene expression analysis for dysregulated immune-related genes between D2-*mdx* vs. B10-*mdx* (**Figure 4**) assessed in juvenile D2-WT and B10-WT hearts. Relative gene expression values normalized to internal *Hprt* transcript levels.

